# Shared and distinct oscillatory fingerprints underlying episodic memory and word retrieval

**DOI:** 10.64898/2026.04.01.715566

**Authors:** Britta U. Westner, Yunzhi Luo, Vitória Piai

## Abstract

Both episodic memory and word retrieval have been linked to power decreases in the alpha and beta oscillatory bands, but these patterns have rarely been related to each other, partly due to a lack of methodological approaches available. In this explorative study, we investigate the similarities and dissimilarities in the oscillatory fingerprints of the retrieval of words and episodes by directly comparing the activity patterns across time, frequency, and space. We acquired electroencephalography (EEG) data of participants performing a language and an episodic memory task based on the same stimulus material. With a newly developed approach, we directly compared the source-reconstructed oscillatory activity using mutual information and a feature-impact analysis. While left temporal and frontal regions showed dissimilarities between the tasks, right-hemispheric parietal regions exhibited similarities. We speculate that this could indicate a homologous function of these regions, potentially sharing less-specific representations between the tasks. We further uncovered a dissociation of the alpha and beta bands regarding the similarity across tasks. While the beta band was dissimilar between word and episodic memory retrieval, the alpha band seemed to contribute to the similarity we observed in right parietal regions. Whether this points to a task-unspecific function of the alpha band or a functional role in the retrieval process of the presumed representations, remains to be determined. In summary, we present an approach to study similarity across tasks using the temporal, spectral, and spatial dimensions of EEG data, and present results of exploring the shared oscillatory fingerprints between episodic memory and word retrieval.

## Introduction

In the conceptual understanding of memory, the declarative memory system has been separated into semantic and episodic memory (Squire, 1992), often studied separately. Semantic memory thereby serves as a repository for factual information about the world (Tulving, 1972), and is directly relevant for language production: Before the ultimate articulation of words, speakers need to retrieve conceptual and lexical information from semantic memory (e.g., Dell, 1986; Levelt et al., 1999). Episodic memory, in turn, refers to the detailed memory of past personal events and experience (Tulving, 2002).

Previous research investigating the neural underpinnings of the retrieval of words and of episodes has employed electrophysiological measurements such as electroencephalography (EEG) and magnetoencephalography (MEG). Of particular interest in this context are brain oscillations: word retrieval in the context of language production (and comprehension) has consistently been associated with oscillatory power decreases in alpha (8-12 Hz) and beta (13-30 Hz) frequency bands (e.g., Hustá et al., 2021; Klaus et al., 2020; Piai et al., 2014, 2015, 2017, 2018, 2020). It has been established that the observed alpha-beta desynchronization effect in word production is not confounded by other processes like attention or motor preparation (for a discussion, see Piai & Zheng, 2019). The alpha-beta effects have been source localized to left inferior parietal lobe, left temporal lobe, and left inferior frontal gyrus (Hustá et al., 2021; Piai et al., 2015; Roos & Piai, 2020), areas associated with semantic memory (Binder et al., 2009) and lexical selection in word production (Indefrey, 2011; Indefrey & Levelt, 2004; Piai & Eikelboom, 2023). Using an auditory distractor paradigm, Piai et al. (2020) experimentally showed that the alpha-beta desynchronization effect is linked to semantic content. Furthermore, a similar alpha-beta power decrease has been identified in language comprehension paradigms (e.g., Rommers et al., 2017). The findings collectively suggest that the observed alpha-beta effects likely reflect a general process of conceptual and lexical retrieval (termed word retrieval henceforth) from semantic memory in language production or comprehension.

Successful encoding and retrieval of episodic memory traces have been linked to increases in synchrony within the theta (3-7 Hz) and gamma (> 40Hz) frequency range, originating from posterior cortical and hippocampal regions (for review, see Nyhus & Curran, 2010). In addition, cortical alpha-beta desynchronization correlates with successful episodic memory formation and retrieval (Hanslmayr et al., 2012, 2016): Alpha-beta power decreases have been repeatedly observed during the encoding of later remembered items compared to later forgotten items, as well as during the correct recognition of encoded items. Prior studies have demonstrated a modulation of the alpha-beta power change by memory retrieval demand, with the accurate recollection of item details eliciting the strongest alpha-beta desynchronization (e.g., Karlsson et al., 2020; Khader & Rösler, 2011; Martín-Buro et al., 2020). Furthermore, alpha-beta power decreases during encoding were predictive of alpha-beta power decreases during retrieval on a trial-by-trial basis (Griffiths et al., 2021). Although memory-related alpha-beta power may not carry stimulus-specific information (Griffiths et al., 2021), the phase of the alpha-beta signal has been shown to code for stimulus identity (Michelmann et al., 2016; Staudigl et al., 2015). Taken together, these studies show that alpha-beta desynchronization is linked to the representation and reinstatement of episodic memory.

Alpha-beta desynchronizations are thus correlated with both language production and episodic memory retrieval processes. Whether this is indicative of a shared retrieval mechanism, however, is unknown. This question is impeded by the fact that there is no clear notion of how the neural underpinnings of the same process in different contexts would exactly be reflected in EEG signals (Francken et al., 2022; Piai et al., 2025). In this study, we set out to explore this question using oscillatory EEG activity. As a first step, we here propose to directly compare the alpha-beta effects between the two retrieval processes to quantify the similarity between the two oscillatory signatures. We characterize similarities and dissimilarities between the two tasks based on the dimensions of time, frequency, and space.

In the present study, we thus compare the oscillatory correlates of word retrieval for language production and episodic memory retrieval within participants. To achieve this, we employ experimental and methodological developments. Experimentally, we combined the context-driven picture naming task, henceforth “the language task”, and an old/new judgment task, henceforth “the memory task” (see Fig. 1 for a depiction of the experiment, following Perrone-Bertolotti et al., 2015). In the language task, participants first listened to incomplete context sentences that were either constrained (i.e., the last word is highly predictable) or unconstrained (i.e., the last word is not predictable), with a picture presented to complete the sentence, following a brief retrieval period. Participants were instructed to name the target picture and memorize the picture in detail. This task is known to produce stronger alpha-beta power decreases during the pre-picture retrieval period in the constrained compared to the unconstrained condition (e.g., Hustá et al., 2021; Piai et al., 2014, 2015, 2020). The subsequent memory task required participants to make old/new judgments on target or foil words, indicating whether they had previously encountered the word as a picture. Following Perrone-Bertolotti et al. (2015), we changed the item presentation modality from pictures in the memory encoding phase (i.e., the language task) to written words in the retrieval phase (i.e., the memory task). The modality change aimed to maximize the recollection-based memory retrieval process in contrast to familiarity-based perceptual judgment (see e.g., Yonelinas et al., 2010 for a discussion on recollection and familiarity). A follow-up question about details of the previously seen picture further prompted memory retrieval. This task elicits stronger alpha-beta power decreases associated with correctly recognized old items compared to correctly rejected new items (Karlsson et al., 2020; Khader & Rösler, 2011; Martín-Buro et al., 2020; Waldhauser et al., 2016).

**Fig. 1:**
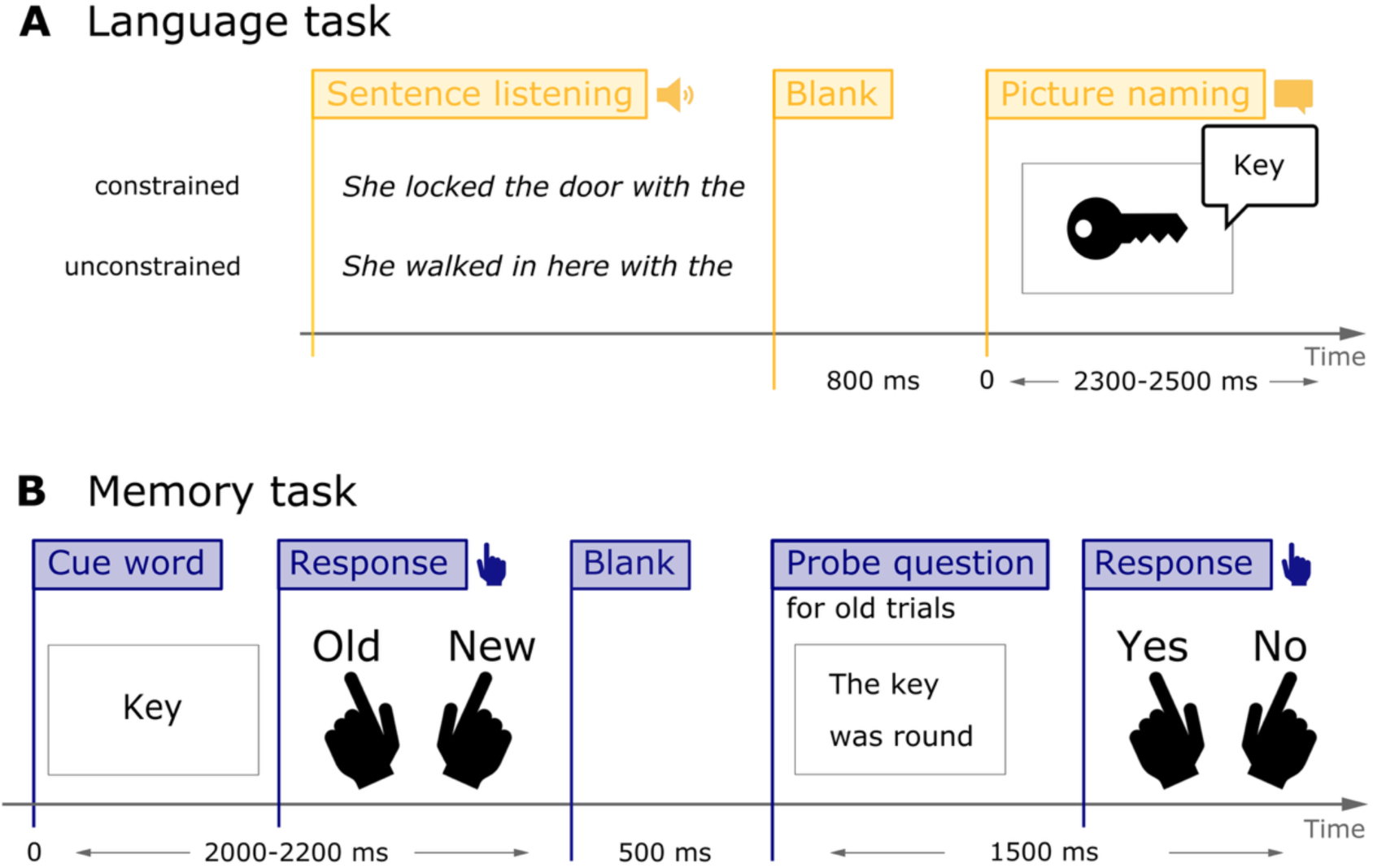
Experiments. **A.** Schematic illustration of the language task. Timings correspond to the events used in the analysis of the data. **B.** Same as A, but for the episodic memory.

The methodological development consisted in comparing the oscillatory effects between tasks, proposing a new analysis approach that quantifies both similarities and dissimilarities across time, frequency, and space. Here, we take an exploratory, not hypothesis-driven approach, describing where and when the two fingerprints are similar and dissimilar. To this end, we assess the similarity of source-reconstructed time-resolved oscillatory activity using mutual information. The similarity analysis is complemented by an impact analysis, revealing which features are similar or dissimilar across the two tasks. If the tasks share a retrieval mechanism, we expect significant similarities in frequency and space (after aligning the time axes between the two tasks based on within-task results, given different time courses following the task demands, see Methods).

We find similarities in right parietal cortex as well as dissimilarities in left temporal and frontal cortex, potentially reflecting the specificity of underlying neural computations to memory and language function. We further report a dissociation between alpha and beta band patterns, raising the possibility that alpha-beta power effects are substantiated by more than one process.

## Methods

### Ethical statement

This study was approved by the Ethics Committee of Faculty of Social Sciences at Radboud University Nijmegen (ECSW-2022-059). All participants provided informed consent before the experiment.

### Participants

Thirty-one healthy Dutch native speakers (7 men; age range: 18-48 with mean age of a subset of 18 participants being 25, exact age missing for 8 participants; normal or correct-to-normal vision; without neurological or language disorders) participated in the experiment, either for course credit or monetary compensation. Data of five participants were excluded from further analyses due to technical issues (n = 1), due to a missing MRI scan (n = 1), because the MRI scan was unusable (n = 2), or because only around 25% of trials were left after artifact rejection due to excessive noise (n = 1). Thus, the data of 26 participants entered the data analysis.

### Materials and design

#### Study design

The study consisted of two tasks: a language task using context-driven picture naming (cf. Piai et al., 2014) and a memory task using an old/new judgment with a shared set of stimuli (following Perrone-Bertolotti et al., 2015). The participants first completed the language task and then the memory task. In the language task, participants first listened to context sentences that were either constrained or unconstrained. Following a brief retrieval period, participants had to name the target picture and memorize the picture in detail. The memory task then required participants to make old/new judgments on target or foil words, thus indicating whether they had previously encountered the word as a picture during the language task. We changed the item presentation modality from pictures in the memory encoding phase (i.e., the language task) to written words in the retrieval phase (i.e., the memory task). The modality change aimed to maximize the recollection-based memory retrieval process in contrast to familiarity-based perceptual judgment (Yonelinas et al., 2010). Between the language and the memory task, there was a short resting state block of 2 min. The data of this block was not used for the analyses reported here.

#### Language task

For the context-driven picture naming task, we selected 106 Dutch sentences with constraining contexts and 106 matching unconstrained sentences, each with the same target word. In the constrained condition, the final word was highly expected based on the context (e.g., “She locked the door with the …” would lead to the target picture KEY). In the unconstrained condition, the context would not prime a specific final word (e.g., “She walked in here with the …” would lead to the target picture KEY). The target pictures were identical in both conditions. Sentences were tested for their cloze probabilities using Welch’s t-test. The cloze probability was significantly higher for the constrained sentences (*M* = 0.89, *SD* = 0.11) as compared to the unconstrained sentences (*M* = 0.01, *SD =* 0.04; *t*(140) = 76.16, *p* < .001). Additionally, the cloze probability for alternative final words in unconstrained sentences was also low (M = 0.22, *SD* = 0.06). All pictures were colored drawings selected from the MultiPic database (Duñabeitia et al., 2018). The context sentences were recorded by a female Dutch native speaker (age: 24). All recordings were segmented and the volume was normalized to 65 dB using Praat (Boersma & Weenink, 2021). The target words were split in two sets and per set, half of the stimuli appeared first in a constrained sentence and then in an unconstrained sentence, and vice versa, presented in a balanced manner across participants. The two stimuli lists within a set were matched for cloze probability and for initial phonemes of target words. Trial order within the two sets was pseudo-randomized using Mix (van Casteren & Davis, 2006) with the constraints that each condition was repeated for a maximum of three trials and that the appearance of the same target picture (in the constrained and unconstrained context) had to be separated by at least ten trials. Participants first completed 8 practice trials with an option to repeat this practice if the procedure was still unclear. The main language task consisted of 212 trials (106 constrained and 106 unconstrained trials), divided into 9 blocks (the last block containing fewer trials than the previous blocks), with self-paced breaks between blocks to minimize fatigue. Each trial started with a fixation cross displayed for 1.5 s before the audio recording of the context sentence was played while the fixation cross remained on the screen. After the recording offset, a blank screen was shown for 0.8 s before participants saw the target picture for a random period jittered between 2.3 to 2.5 s, after which a screen was displayed for 2 s which signaled to participants to blink if necessary. For a visual representation of the key aspects of the language experiment, see Fig. 1A.

#### Memory task

For the episodic memory retrieval task, we paired the 106 target words used in the language task with 106 foil words. The target and foil words were matched for the most relevant word properties that would affect memory recall performance: animacy, size of the item, arousal, and word length (Madan, 2021). The selected target and foil words had exactly the same distributions for animacy (animate: 33 items; inanimate: 73 items) and size (51 small items; 55 large items) as categorized by the “shoebox size judgement test” (i.e., whether the item would fit into a shoebox). Target and foil words were comparable in terms of arousal level (target: *M =* 3.83*, SD =* 0.89, range: 2.03-6.22; foil: *M =* 3.80*, SD =* 0.7; *t*(195) = 0.21, *p* = 0.83), as rated in the normed Dutch word database by Moors et al. (2013). There was no difference in word length measured by number of letters (target: *M* = 4.83, *SD* = 1.31; foil: *M* = 4.83, *SD* = 1.47; *t*(207) = 0, *p* = 1). Lexical frequency retrieved from the SUBTLEX-NL database (Keuleers et al., 2010) was also comparable between target (*M* = 4.25, *SD* = 0.57) and foil words (*M* = 4.20, *SD* = 0.47; *t*(202) = 0.78, *p* = 0.44). To additionally measure the depth of episodic memory retrieval in the memory task, each trial was followed by a sentence describing a detail of the language-task picture (e.g., for target picture KEY the description was “The key was round in shape.”). This description could be either accurate or inaccurate. Before the study, we administered an online test to ensure that people generally agreed on the accuracy for the description of each picture. Fifteen Dutch native speakers (mean age of 20, 2 male) participated in this online test but not in the main experiment. The descriptions passed the selection criterion if at least 12 out of the 15 participants agreed on the correctness of the description, which could be either correct or incorrect. Four descriptions had to be revised until all passed the criterion. Trials for the memory task were pseudo-randomized. A unique list was generated for each participant. The same condition (i.e., whether the item was old or new) never appeared more than three times on subsequent trials. Similar to the language task, participants did 8 practice trials and could choose to repeat the practice. The memory task contained 212 trials (106 old and 106 new) that were split into 9 blocks (with the last block containing fewer trials than the previous blocks). Each trial began with a fixation cross for 1.5 s before the cue word was presented for a random duration jittered between 2 and 2.2 s. Participants were tasked to indicate as quickly as possible whether they had seen the word as a picture in the previous language task. Participants responded by pressing a button with either the index or middle finger of their left hand. The assignment of the two fingers to “yes” or “no” was counterbalanced across participants. If participants answered “yes” (indicating an old word) correctly, the corresponding written description would be shown on the screen after a 0.5 s blank screen. The description was presented until a button response was made. Lastly, the blinking screen was displayed for 1.5 s. A depiction of the key aspects of the memory experiment is shown in Fig. 1B.

### Data acquisition

All auditory and visual stimuli were presented with the Presentation software (Neurobehavioral Systems Inc., Berkeley, United States). EEG was recorded using an Acticap system with 64 electrodes and a BrainAmps DC amplifier (Brain Products GmbH, Gilching, Germany) with a sampling rate of 500 Hz and an online band-pass filter of 0.016-125 Hz. The online reference was the left mastoid. In addition, we used seven passive electrodes to record the vertical and horizontal electrooculogram as well as the mouth electromyogram, for which we placed electrodes on the right upper and lower side of the orbicularis muscle. To ensure optimal recording quality, impedances of all electrodes were brought to below 15 kΩ. After the EEG recording, the cap placement was captured using a 3D scanner to be used for MRI-coregistration and electrode positions in forward modelling.

T1-weighted structural MRIs were acquired on a 3T MAGNETOM Prisma or PrismaFit scanner (Siemens AG, Healthcare Sector, Erlangen, Germany) using a 32-channel head coil. Participants lay in the supine position during the acquisition. The scan was acquired in the sagittal orientation using a 3D MPRAGE sequence with the following parameters: a repetition time of 2.3 s, an inversion time of 1.1 s, and an echo time of 3 ms with an 8° flip angle. The field-of-view was 256 × 225 × 192 mm and we used a 1-mm isotropic resolution. The acquisition time was 5 min 21 sec.

EEG and behavioral data will be available upon journal publication.

### Behavioral data analysis

#### Language task

During the language task, participants’ verbal picture naming responses were recorded for later analyses of accuracy and response time (RT), to compare to previously published results. Audio recordings began at the onset of the target picture and lasted for 3.5 seconds. Only correct responses or close synonyms (e.g., “bird nest” instead of “nest”) uttered after picture onset were counted as correct responses. Incorrect responses, responses with hesitation, self-correction, or determiners, and early responses were counted as incorrect and removed from all subsequent RT and EEG analyses. RTs were manually marked for each trial in Praat (Boersma & Weenink, 2021) blinded for conditions. Participants’ mean RTs were compared between conditions using a dependent samples t-test in MATLAB (version R2023b; The MathWorks Inc., 2023).

#### Memory task

In the memory task, participants’ button press responses were automatically coded as correct or incorrect during the experiment. We then manually marked any items as invalid for which the participant had produced a label other than the target picture name during the language task (e.g., if a participant produced ‘mug’ for the intended target *cup*, the item ‘cup’ in the memory task was marked as invalid). Accuracy was computed per condition based on the correct and incorrect/invalid trials. Only correct, valid responses were included in subsequent EEG analyses.

### EEG data analysis

EEG data analyses were performed using FieldTrip (version hash: 7ba0239; Oostenveld et al., 2011) and custom code in MATLAB (version R2023b; The MathWorks Inc., 2023). Some of the colormaps for data visualization were created using the cmocean package for MATLAB (Thyng, 2020). The analysis code for the project will be available upon journal publication.

#### Forward modelling for EEG source reconstruction

Forward models for source reconstruction were computed based on individual magnetic resonance images (MRIs). If necessary, MRIs were bias corrected using SPM12 (Wellcome Centre for Human Imaging) prior to further processing. MRIs were then resliced and segmented into brain, skull, and scalp. Meshes were prepared for all three tissue types, and a boundary element model (BEM) volume conductor was created using the “dipoli” method (Oostendorp & Van Oosterom, 1989). We identified electrodes on a 3D scan of the participant wearing the EEG cap to get individual electrode locations. This 3D scan was also used to semi-automatically align the coordinate systems of the MRI and the EEG electrodes using facial features from the MRI scan and the 3D scan (GitHub repository: https://github.com/britta-wstnr/source_recon_langdysfun). Source models were constructed based on an 8 mm template grid in MNI space and warped to individual space. Based on all this information, forward model solutions were computed using FieldTrip.

#### Preprocessing

The raw EEG data was epoched into trials. For the language task, data was epoched from −1.8 to 1.0 s relative to picture onset (cf. Fig. 1A) and for the memory task, data was epoched from −0.5 to 2.3 s relative to the cue word onset (cf. Fig. 1B). Only correct, valid trials were kept for further analysis. Corrupted or overly noisy channels were removed and equalized across tasks (63.85 channels remaining on average across both tasks, *SD* = 0.37). Independent component analysis (ICA) was performed on the appended data of both tasks, and eye movement and heartbeat components were removed from the data (2.27 components removed per participant on average, *SD =* 0.67). The data was re-referenced to an average reference. The epochs were demeaned but not filtered as subsequent analyses were exclusively done in the frequency domain. Noisy trials were rejected using a threshold criterion of 100 µV. This automatic trial rejection was subsequently manually inspected. After trial rejection, there were on average 74.96 trials in the constrained condition (*SD =* 15.64) and 76.15 trials in the unconstrained condition (*SD =* 16.12) of the language task. There was no significant difference in the number of trials between the conditions for the language task (*t*(50) = 0.27, *p* = 0.788). In the memory task, there were on average 98.85 trials in the new condition (*SD =* 3.41) and 80.62 trials in the old condition (*SD =* 18.89). There was no significant difference in number of trials between the conditions for the memory task (*t*(50) = 0.72, *p* = 0.472). The discrepancy between the language and memory task in terms of removed trials is rooted in the word production aspect of the language task. While we relied on pre-utterance data for our ultimate analyses, we still had to include some of the signal after picture presentation to allow for time-frequency decomposition. Since this signal inadvertently gets smeared into the pre-utterance window, we included it in the cleaning procedure, which in turn leads to a higher number of trials rejected in the language than in the memory task.

#### Within task analysis

The channel-level data was transformed to the frequency domain using a time-frequency approach based on the Fast Fourier transform. Frequencies from 4 to 30 Hz were estimated with a frequency resolution of 1 Hz. The sliding time window with a Hanning taper was adapted to the frequencies, spanning 5 cycles. The window was centered every 10 ms. A dependent samples t-Test was run for each task, testing for the difference between conditions. We tested at an alpha level of 0.05 and only one-sided, since a decrease in alpha/beta power was expected based on both the language (for review, see Piai & Zheng, 2019) and memory literature (for review, see Fellner & Hanslmayr, 2017). To correct for multiple comparisons, we employed the Max-T Statistic embedded in a Monte Carlo permutation scheme with 10,000 permutations (Nichols & Holmes, 2002; Westfall & Young, 1993). For plotting only, the two conditions per task were subtracted and the resulting difference was divided by the average of the two conditions. Based on time-frequency maps masked by the multiple-comparison-controlled significance, a time-frequency window was identified for source reconstruction. This was done jointly for both tasks, such that the window length and frequency content were identical for both tasks and captured the condition effect for both. The window was 0.5 s long and spanned 6-10 Hz. For the memory task, the window was centered on the strongest difference between the conditions at 1.15 s after cue onset (cf. Fig. 4B). For the language task, the window was centered such that it did not extend into the post-picture interval, i.e., at −0.75 s relative to picture onset (cf. Fig. 4A). These windows were used as input to a Dynamic Imaging of Coherent Sources (DICS) beamformer (Gross et al., 2001) with unit-noise-gain (Borgiotti & Kaplan, 1979). The CSD matrices were computed across conditions but per task (common spatial filters) and were regularized with 15% of global electrode power. CSD matrices were inverted using a truncated pseudo inverse (Westner et al., 2022). The resulting task-specific beamformer weights were then separately applied to both conditions per task. The resulting source images were then subtracted from each other and then divided by the average of the two conditions. The results were then masked to only show the 5% source points with the strongest power decreases.

### EEG similarity analysis

The similarity analysis of source-reconstructed time-frequency data was performed to compare the data of the two tasks. Using mutual information, we quantified the similarity within and across tasks. We then deployed an impact analysis to identify which features of the data were similar and dissimilar across the tasks.

For the similarity analysis, only the trials of the constrained condition in the language task and the old condition in the memory task were used. This was motivated by the fact that only in these conditions, a retrieval of a word before picture presentation (language task) or the retrieval of an episode after cue presentation (memory task) was possible. Due to the already discussed discrepancy in trial numbers between the two tasks, we subsampled the task with the higher number of trials if the difference in trial numbers was higher than 20%. This was done to prevent the data covariance matrix for the beamformer to be skewed towards one task while keeping as many trials as possible. After subsampling, there were on average 74.96 trials left for the language task (*SD =* 15.64) and 80.62 trials for the memory task (*SD* = 18.89). There was no significant difference between the numbers of trials between the two tasks (*t*(50) = 1.18, *p* = 0.245). The data was then source-reconstructed into atlas parcels. We used the Brainnetome atlas (Fan et al., 2016) as shipped with FieldTrip. Deep structures (i.e., structures that are subcortical and do not lie in the cortex hull or medial wall) were manually excluded (114 parcels in total), and small adjacent parcels were merged with each other to prevent parcels with less than 3 grid points. After these two steps, 126 out of 246 parcels remained across both hemispheres. For each parcel, we then computed the strongest component of the leadfield via singular value decomposition (Backus et al., 2016) and used this information to obtain the representative source-reconstructed activity of a parcel via a linearly constrained minimum variance (LCMV) beamformer (Van Veen et al., 1997) with the same specifications as the DICS beamformer reported above. The time windows for the two tasks were chosen to be 1 s long and to represent the retrieval windows for language and memory, respectively. For the language task, we chose the 1 second leading up to the picture presentation (cf. Fig 1A, also see Fig 4A). For the memory task, we chose a time window from 0.7 to 1.7 s after word cue onset (cf. Fig. 1B, also see Fig. 4B).

The spatial filter was computed across both tasks, as in the subsequent analyses the two tasks were compared against each other. The single trial source data was then transformed to the frequency domain for the chosen time windows with the same approach as reported above, except that the sliding time window was now centered only every 25 ms to reduce computational load on the subsequent analyses. The obtained time-frequency-resolved data was log-transformed and saved for both tasks separately.

The similarity analysis was done in Python, using custom code (link to repository will be available upon journal publication). The basis of this similarity analysis is the metric of mutual information. To acquire a “ceiling” of similarity, we first computed similarity *within* tasks per participant. After halving the data, the algorithm randomly picks two sets of *n* trials from each half, averages these *n* trials, and computes the mutual information between the two sets (see Fig. 2A for an illustration of the process). The computation of the mutual information was implemented following Cohen (2014). The *n* is then iteratively increased from one trial to the full *N*, providing a similarity curve as a function of trials in the comparison. The full procedure is repeated 100 times and results are averaged across the 100 outcomes. The mutual information curve was used for benchmarking, ensuring that the similarity reaches a plateau when approaching the full set of trials. After computing mutual information as a measure of similarity within tasks, we repeated the approach picking one set of the trials from the language task and one set of trials from the memory task to see how overall similarity between the tasks compares to within-task similarity.

**Fig. 2:**
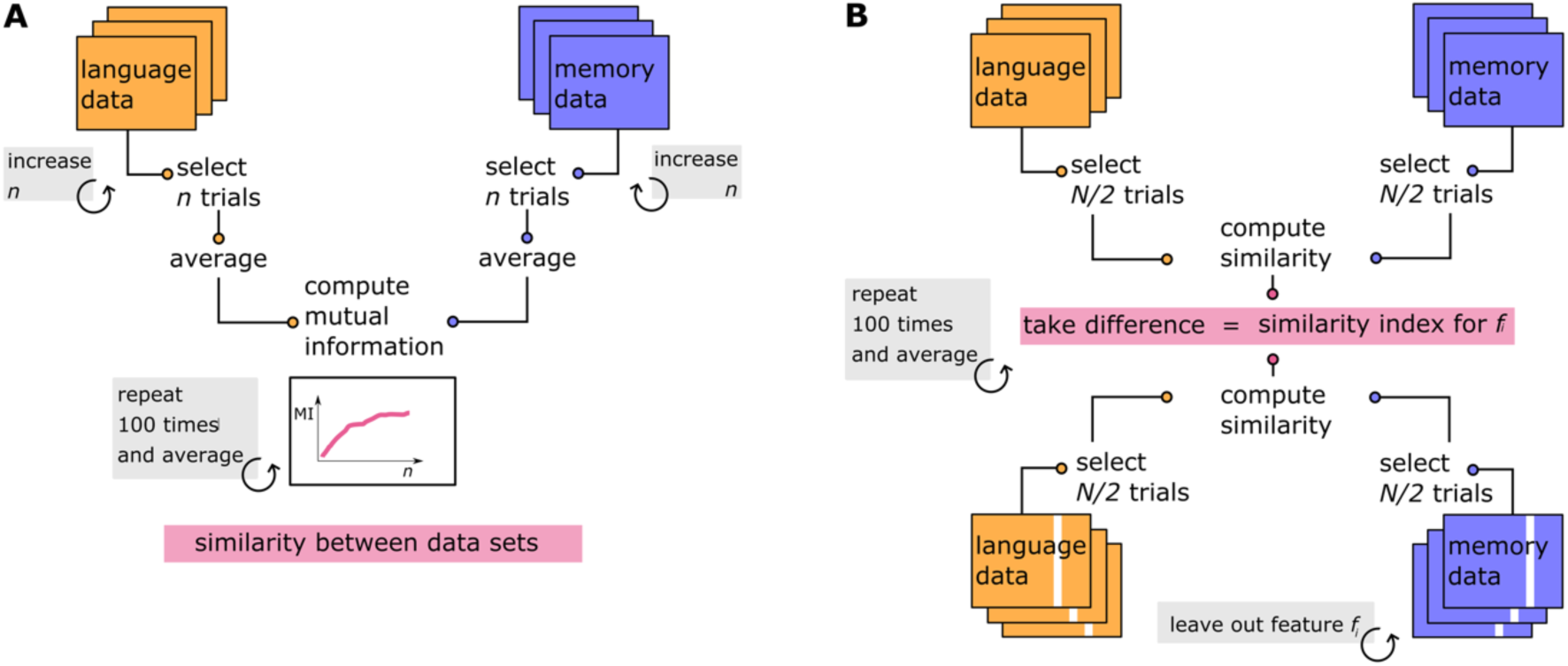
Similarity analysis. **A.** Computation scheme for between-task similarity analysis. **B.** Impact analysis, estimating the similarity of certain features (e.g., a specific frequency) between tasks.

Next, we quantified the impact that single features have on the similarity. In this context, features are the time points, frequencies, and parcels that span our data. Thus, the question for this analysis was, for example: how big is the impact of 10 Hz on overall similarity? This translates to: how similar is the data at 10 Hz? To answer this question, we took inspiration from variable importance measures in decision trees and forests (Breiman, 2001; Breiman et al., 2017). This analysis was done per participant. Leaving one feature out, the mutual information between two the datasets was recomputed (using a random sample of half of the data, repeating this procedure 100 times to decrease the impact of potential outliers in the drawn sets of trials) and then compared to the similarity when including that feature (see Fig. 2B for an illustration of the process). If the similarity decreases as compared to the full dataset, the removed feature was more similar across datasets; conversely, if the similarity increases, the removed feature was more dissimilar. For ease of interpretation, we multiplied the resulting change in mutual information by −1 such that the measure (which we call similarity index in this study) increases if the feature is similar and decreases if it is dissimilar. The described procedure was then repeated for all features in the dataset (i.e., 27 frequencies, 41 time points, and 126 parcels).

The resulting similarity indices were then tested against zero across participants (null hypothesis: leaving out this feature has no impact on similarity) using a cluster permutation test for each feature type, that is, frequency, time, and parcels. The cluster permutation test was used as implemented in MNE-Python (Gramfort et al., 2013; version 1.7.0: Larson et al., 2024). For frequency and time, a linear neighbouring structure was assumed, while for parcels, neighbours were identified based on the distance of the centers of mass among parcels.

## Results

### Behavioral results

#### Language task

Participants were more accurate at naming pictures in the constrained than in the unconstrained condition (Fig. 3A). In the constrained condition, participants were on average 96.26% correct (*SD* = 3.62), while they were on average 93.32% correct in the unconstrained condition (*SD* = 3.49). There was a significant difference in accuracy between the conditions (*t*(25) = 3.49, *p* = 0.002). For the language task, we also reproduced the well-established RT effect (Fig. 3C): The participants were significantly faster in naming the picture in the constrained (*M =* 0.60 s, *SD* = 0.11) than in the unconstrained condition (*M =* 0.83 s, *SD* =0.07; *t*(25) = −18.83, *p* < 0.001). Note that this subset of the behavioral data was already reported in Chupina et al. (2025).

**Fig. 3:**
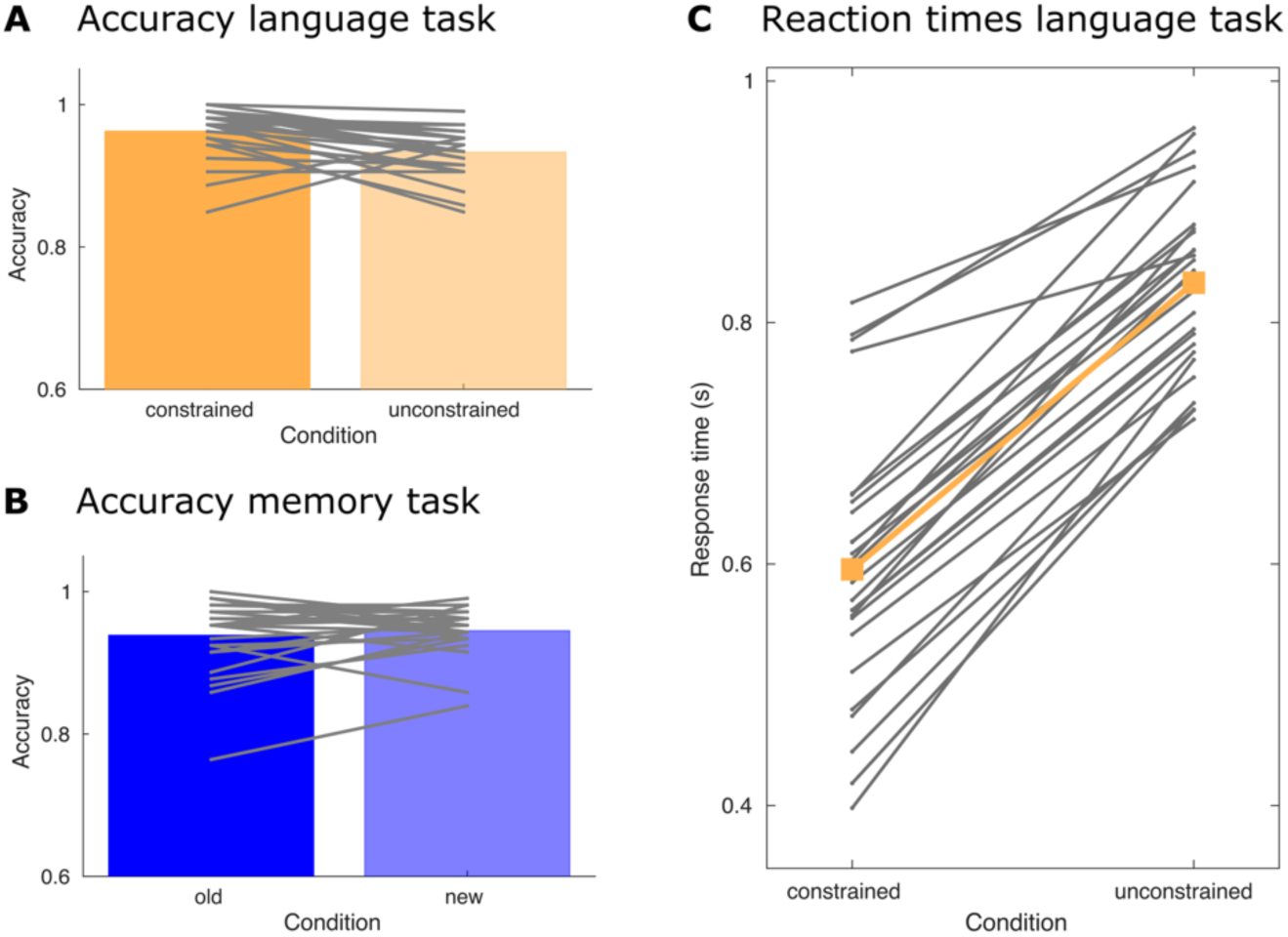
Behavioral results. **A.** Participant-level (lines) and group-level (bars) averages for task accuracy in the language task **B.** Same as A, but for the memory task. **C.** Participant-level (grey lines) and group-level (orange line) averages of reaction times in the language task.

#### Memory task

The participants correctly identified old items with an accuracy of 93.87% (*SD* = 0.05) and new items with an accuracy of 94.52% (*SD* = 0.03). There was no significant difference in the performance between the two conditions (*t*(25) = −0.76, *p* = 0.457; see Fig. 3B).

### Within-task EEG results

#### Language task

The electrode-level time-frequency representation of the contrast between constrained and unconstrained sentences revealed the expected alpha-beta desynchronization prior to target picture presentation (Fig. 4A). The test across time, frequency, and electrodes revealed only a small portion of this alpha-beta desynchronization to be significantly different between the conditions (Fig. 4C; corrected for multiple comparisons using a Max T-Statistic and Monte Carlo permutation, cf. Methods section). This effect is concentrated at the boundary of the theta and alpha band (6-7 Hz) and situated rather late, coinciding with the presentation of the target picture. The source reconstruction of the alpha power decrease during the 0.5 s preceding target picture onset localizes mainly to left temporal and central areas (Fig. 4E). Scattered right-hemispheric regions are indicated as well, e.g., lateral occipital, parietal, and frontal regions. Note that the chosen time-frequency window for the source reconstruction was informed not only by the language, but also by the memory task.

**Fig. 4:**
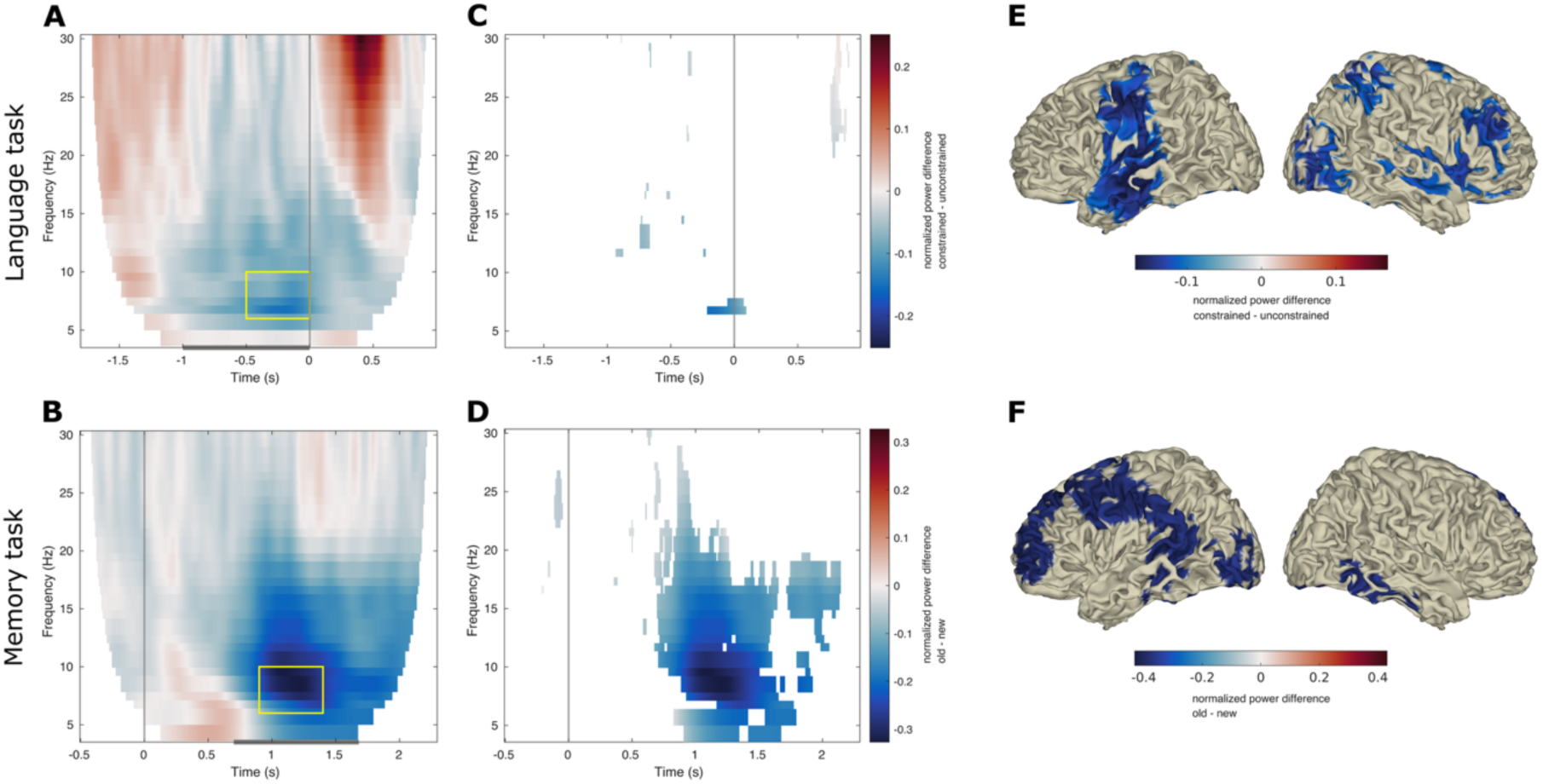
Within-task results. **A.** unmasked time-frequency plots for language (top) and memory task (bottom). For the language task, the constrained condition was compared to the unconstrained condition. For the memory task, the new and old items were compared. Note the different time axes (0 marked with a vertical line). 0 corresponds to naming cue onset (picture) in the language task and the memory probe (word) in the memory task. The yellow windows mark the time windows used for source reconstruction (C). The grey bar on the x-axis marks the time windows used for the similarity analysis. **B.** Results from within-task permutation test for language (top) and memory task (bottom). **C.** Source reconstruction results from a DICS beamformer based on the yellow boxes in (A). The frequency range was kept the same across tasks (6-10 Hz), and both windows are 500 ms long (language: −0.5 to 0 s; memory: 0.9 to 1.4 s).

#### Memory task

The time-frequency representation of the memory task shows a pronounced alpha-beta power decrease centered around 1.25 seconds after target word onset and spanning from alpha far into the beta spectrum (Fig. 4B). Testing these electrode-level data revealed a significant difference between the conditions, substantiated between 0.7 and 2 s after cue word presentation and spanning almost the whole analyzed frequency range (Fig. 4D). The source reconstruction of the alpha decrease (6-10 Hz) between 0.9 and 1.4 s localizes mainly to frontal, central and temporal regions in the left hemisphere as well as inferior temporal regions in the right hemisphere (Fig. 4F).

#### Identification of task-specific time windows

For the similarity analysis, we needed to identify 1 second time windows per task which contain word and episode retrieval, respectively. We based these time windows on the obtained within-task results as well as on the literature. Hereby, the comparably rather weak alpha-beta desynchronization in the language task was surprising (cf. Hustá et al., 2021; Klaus et al., 2020; Piai et al., 2014, 2017, 2020; Roos & Piai, 2020 for previously published effects). We therefore based the language task time window on this literature as well as the raw alpha-beta power decreases (Fig. 4A) instead of on the test results (Fig. 4C). To minimize the leakage of post-picture presentation effects, we did not expand the time window beyond the timepoint of picture presentation. All these considerations led to a time window spanning the second before picture presentation for the language task. For the memory task, we were strongly guided by the within-task results. After comparison with the literature (Burgess & Gruzelier, 2000; Düzel et al., 2003; Karlsson et al., 2020; Spitzer et al., 2008) we chose a time window ranging from 0.75 to 1.75 s after cue word onset. The time windows are marked in the within-task results as grey bars (Fig. 4A and B).

### Similarity analysis

We first computed the similarity within tasks to serve as an upper boundary. This was done per participant and repeatedly, increasing trials. Figure 5 shows the within-task similarity in yellow for the language and blue for the memory task, averaged over participants. Note that with increasing trial numbers, the number of subjects in the average similarity decreases (see grey density at the bottom of the figure), meaning that the curves become less generalizable beyond a certain point as only few participants supplied enough trials (above 80; note that the data is split into two halves for the comparison). This first computation serves as a sanity check of sorts: First, as expected, the similarity increases with increasing trial numbers. The mutual information seems to plateau between 30 and 40 trials per average (before there is an increase again which, as argued above, is not generalizable due to low subject count). Interestingly, the within-task mutual information is highly similar for both tasks. We then computed the between-task similarity in the same manner, now taking one average of the language task and one of the memory task. As expected, the between-task similarity is lower than the within-task similarity. However, the mutual information curve has roughly the same shape and is in the same order of magnitude, suggesting shared features between the two data sets.

**Fig. 5:**
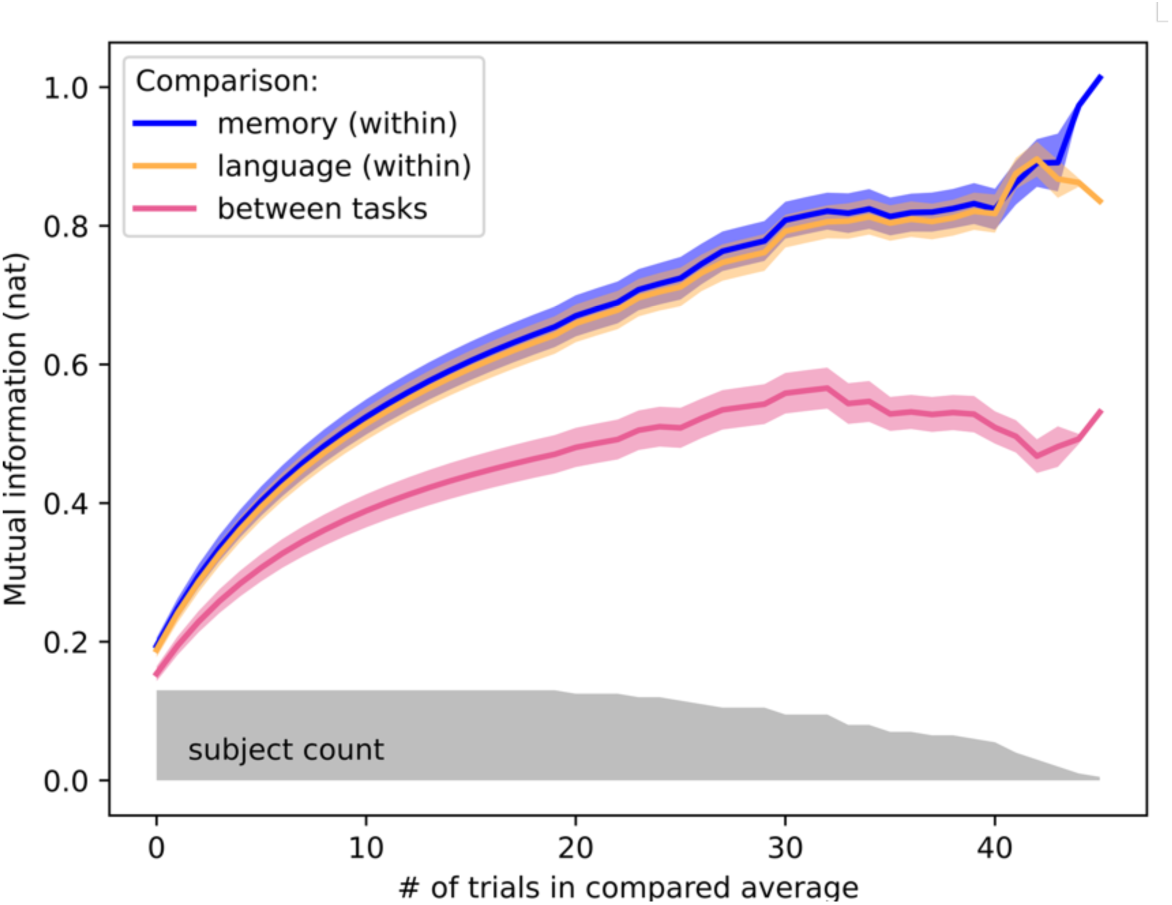
Similarity within and between tasks. Similarity (mutual information) between data averages as a function of number of trials in the average. All comparisons are done within subject and then averaged across subjects. Since subjects have different trial numbers, higher numbers of trials in the average are represented by less subjects. This is indicated by the grey subject count distribution (range: full N=26 on the left, N=1 on the right). Blue curve: Mutual information for the within task comparison for the memory task, yellow curve: within task comparison for the language task. Pink curve: between task comparison, average of comparing memory to language and vice-versa. Shaded areas represent the SEM.

To find out which features across time, frequency, and space are shared between the two data sets (and which are dissimilar), we next did an impact analysis. Leaving only one feature out at a time, we recomputed the similarity between the two tasks and compared the result with the original mutual information when all features were present (cf. Fig. 2B for a schematic of the analysis). We report the results of this impact analysis in Figure 6.

**Fig. 6:**
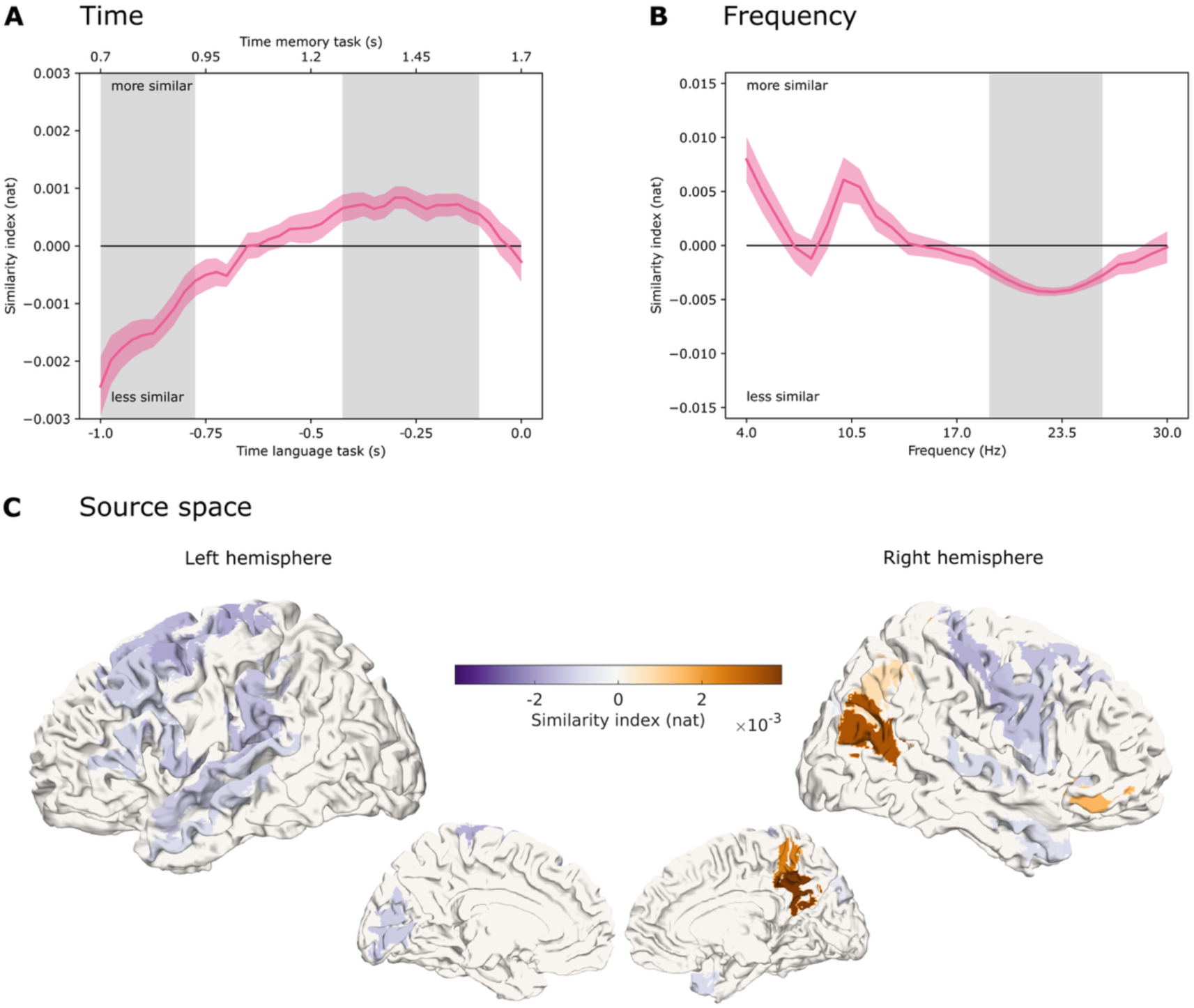
Impact analysis (analysis of feature similarity). **A.** Impact analysis between tasks. The values now represent whether a certain time point is similar (positive) or dissimilar (negative) when comparing between the memory and the language task. Note again, that the time points differ between the tasks (two x axes). The grey areas signify clusters of the permutation test. **B.** Same as B, but for frequency instead of time. **C.** Impact analysis between tasks for brain regions. Shown are brain regions that formed clusters in the cluster permutation test. Purple hues show dissimilar regions, brown hues show similar regions.

#### Time

We find clusters of dissimilar and similar time points (cf. Fig. 6A), with earlier timepoints in the timeseries being dissimilar and later timepoints being similar. Referencing the within-task results of the language task (cf. Fig. 4A), the variance seems to be least towards the end of our time window (cf. statistical results in Fig. 4C). On the other hand, the earlier timepoints show an increase in alpha power for the memory task (cf. Fig. 4B), presumably evoked by the stimulus presentation. We can thus speculate that the late increase in similarity could index the capturing of the retrieval process in both datasets. However, the time dimension is arguably the least suited for this kind of similarity analysis. Within tasks, we can expect variations in exact timing of sub-processes (Chupina et al., 2025) and between tasks, we expect even less comparability. While we carefully decided for the time windows per task such that they reflect the retrieval process, comparing the tasks timepoint-by-timepoint (as done when removing specific timepoints from the analysis) is potentially comparing mis-aligned processes.

#### Frequency

We find high similarity in the alpha range, and dissimilarity in the beta range (Fig. 6B). While the alpha similarity does not substantiate a cluster, despite descriptively showing high similarity, the curve is significantly different from zero with a dissimilar cluster in the beta range from 19 to 26 Hz.

#### Source space

In source space, there are several similar and dissimilar parcels (cf. Fig. 6C, Supplementary Table 1 and Supplementary Figure 1). Similar regions are concentrated in the right hemisphere, in parietal and inferior frontal parcels. Dissimilarity is seen in frontal and temporal regions in both hemispheres. Hereby, the motor regions can serve as another “sanity check” for the analysis: we expect some reactions (i.e., finger movements of the left hand) to fall within the analyzed time window of the memory task, while the language task time window may contain preparatory articulatory activity. We thus expected these regions to show topographical differences, which is shown by the impact analysis results over motor regions.

Lastly, we visualized the time-frequency representations for the similar regions and some of the dissimilar regions. To this end, we averaged the time-frequency representations across all similar or dissimilar parcels within the same region. Figure 7A depicts the time-frequency representations of three similar regions, right precuneus, right inferior parietal lobule, and right orbitofrontal gyrus, shown as the difference between language and memory task, averaged across participants. The color bar limits are scaled to the limits of the individual representations for language and memory. In posterior regions, early alpha band activity seems to be consistently higher for the language task. Note that this is not directly comparable to the within-task results reported in Figure 4, as we are only looking at data from the retrieval condition here (constrained condition for the language task and old condition for the memory task), not at differences between conditions. The difference between the tasks becomes smaller at later timepoints, especially for the upper and lower bound of the alpha band. There is comparably less difference in the beta band. For the orbital frontal gyrus, the comparison shows a slightly higher power across many frequencies (alpha and beta band) and the whole time window for the language as compared to the memory task. In Figure 7B, we report some of the dissimilar regions in the left hemisphere: superior and middle temporal gyrus and inferior frontal gyrus (see Supplementary Figure 2 for additional regions). Being typical language regions, they all show a comparably higher alpha and beta power in the language than the memory task. Interestingly, inferior temporal gyrus, which might be implicated in the memory task (cf. Fig. 4F) showed neither similarity nor dissimilarity between the tasks.

**Fig. 7:**
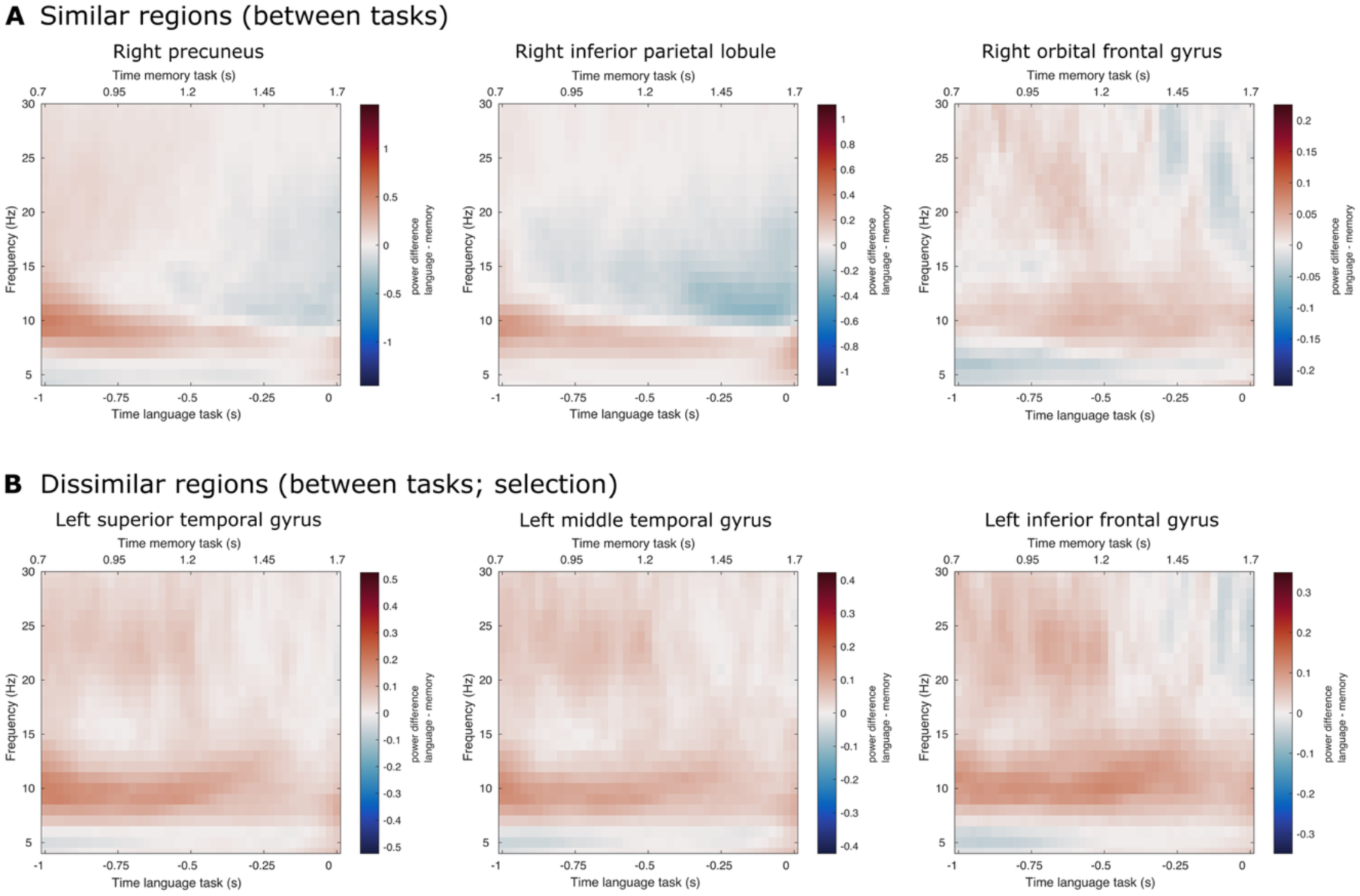
Time-frequency task contrasts in regions of interest. Contrasting time-resolved oscillatory power between the language and memory task (retrieval conditions only; *constrained* for the language task and *old* for the memory task). Red colors represent a higher absolute power in the language task, blue colors represent a higher absolute power in the memory task. Color limits are based on the maximum absolute power across both tasks in the given region of interest. Note the different time axes for the memory task (top) and the language task (bottom). **A** Regions that belong to a similar cluster. **B** Regions that belong to a dissimilar cluster.

## Discussion

In this study, we assessed the similarities and dissimilarities across the source-reconstructed oscillatory fingerprints of word and episodic memory retrieval. We acquired EEG data while the participants completed a language and a memory task with shared stimulus material. We analyzed the data using a newly developed similarity analysis based on mutual information. The subsequent impact analysis revealed similarities between the two tasks especially in right parietal regions and within the alpha band.

### Within-task results

To establish task-specific retrieval effects, we first analyzed the behavioral and electrophysiological data within the tasks. For both the language and the memory task, the behavioral data follow previously published patterns. The time-frequency retrieval effects for the memory task (old/new contrast) show the expected alpha-beta decreases. The language task shows a weaker oscillatory effect for word retrieval (contrast of constrained versus unconstrained condition) than previously reported but the overall time-frequency pattern and behavioral results point to a retrieval effect in the data. In summary, the within-task results confirm the presence of retrieval in both tasks. For further analyses, we then looked at the retrieval conditions only (constrained condition of the language task and old condition of the memory task), which means that further analyses are based on absolute power (and not power differences between conditions).

### Assessing similarity in electrophysiological data

In this paper, we present a new method to investigate similarities and dissimilarities in oscillatory fingerprints between datasets. The method is based on mutual information and provides an impact analysis for all features (e.g., space, time, and frequency). This allows for testing the impact against zero, which provides significance testing for similar *and* dissimilar features. Here, we chose to use a two-sided cluster-based permutation test, which provides us with negative (dissimilar) and positive (similar) clusters that substantiate the rejection of the null hypothesis that there is no impact of a given feature on mutual information. We combined these results with a visualization of the raw power effects between tasks.

Crucially, mutual information is an entropy-based measure that quantifies the amount of information that is shared between two variables and is thus based on the amount of information present within each variable. If two variables share the absence of a signal, that would therefore not lead to high mutual information, even though the variables are “similar”. In the following, we will first argue that this method delivers expected outcomes (e.g., when it comes to diverging task demands) and then further consider the retrieval-based results we obtained. Lastly, we discuss shortcomings of the approach when it comes to the timing of the observed effects.

### Dissimilarities across tasks match diverging task demands

The impact analysis shows dissimilarities in especially right motor regions (see Fig. 6C and supplementary Figure 1), which matches with the different task demands: the time window used for the analysis of the memory task contained the reactions of the participants (left-hand button press). The language task time window may further contain bilateral preparatory motor activity, which matches with the dissimilarities in the wider motor areas in both hemispheres. Furthermore, the beginning of the language task time window may contain activity stemming from the offset of the auditory presentation of the sentence, which ended at −800 ms relative to picture onset at 0 ms. This matches the dissimilarities we see in superior temporal gyrus and wider auditory regions.

### No shared fingerprints in temporal regions

In the typical language and memory regions in the left hemisphere, memory and word retrieval did not share oscillatory fingerprints. In left-hemispheric temporal and frontal regions, the algorithm uncovered dissimilarities between the two tasks (Fig. 6C; see Supplementary Table 1 for details). Looking at the time-frequency representations in these areas shows that the difference in the power time course between the two tasks is largest in the alpha range (Fig. 7B), with more alpha power being present in the language task. Indeed, alpha power across all electrodes was higher for both conditions in the language than in the memory task (Supplementary Figure 3). This also aligns with the relatively smaller alpha power decrease between conditions for the language task (Fig. 4E) as compared to the memory task (Fig. 4F). The order of the tasks was always the same across participants with the language task presented first (given that the language paradigm doubled as the encoding phase for the memory task), which means that a simple fatigue effect leading to higher alpha power would be expected to emerge in the memory task, contrary to what we find. Overall higher alpha power in the language task does not drive all our dissimilarity results either: some regions have a higher alpha power in the memory task (cf. Supplementary Figure 2), that is, the reverse pattern. Notably, our impact analysis based on mutual information goes beyond a direct comparison of power. In the following paragraphs, we explore why similarities and dissimilarities might emerge between the tasks.

### Similarities in right parietal regions: A homologue area?

We find similarities between the two tasks in right precuneus and right inferior parietal lobule (see Supplementary Table 1 for details). The parietal lobe, including the temporo-parietal junction, has been associated with episodic memory retrieval (e.g., Rugg & Vilberg, 2013; Wagner et al., 2005) and word retrieval in language (Binder, 2015; Binder et al., 2009; Chang et al., 2017; Rugg & Vilberg, 2013). Notably, the literature consistently reports a left-hemisphere dominance for word retrieval (Riès et al., 2016).

In old/new memory paradigms, the lateralization of the alpha/beta desynchronization effect seems to depend on the used stimulus material: While language stimuli elicit a left-lateralized effect (e.g., Düzel et al., 2003; Spitzer et al., 2008), faces for example lead to a lateralization on the right (Burgess & Gruzelier, 2000). This has led to speculation that the parietal alpha-beta desynchronization reflects a process specific to memory retrieval, possibly the reactivation of sensory features of a memory trace (Hanslmayr et al., 2012). This has further been solidified by reports of content-specific reinstatement of temporal patterns in electrophysiological signals (Jafarpour et al., 2014), including alpha (Michelmann et al., 2016) and beta activity (Staudigl et al., 2015). Remarkably, Michelmann et al. (2016) showed that for their visual condition, alpha phase similarity between encoding and retrieval is localized to *right* superior lateral and medial parietal cortex.

Here, we compared a word and a memory retrieval task, with the language task doubling as the memory encoding phase. More precisely, we compared the pre-picture interval of the language task with the memory retrieval phase cued by a word presentation. The material the participants are instructed to remember, however, are the pictures they named in the language task, thus the memory retrieval phase in the second experiment would probably reinstate the visual experience of these pictures. We thus do not strictly compare the encoding and retrieval phase as the papers referenced above do. Yet, following the idea that memory alpha-beta desynchronization effects could reflect the reinstatement of the memory trace, the memory retrieval phase in the second experiment would probably reinstate the visual experience of these pictures, which would fit the lateralization of our raw power effects to the right hemisphere.

Word retrieval, by contrast, would lead to an expectation of activity in the left-hemispheric areas. In our experiment, however, the time-frequency representations of left-hemispheric areas are neither part of a similar nor dissimilar cluster. However, it has been reported that the right hemisphere is also involved in verbal processing, with homologue areas showing an – albeit weaker – contribution in language tasks (Just et al., 1996; Martin et al., 2022), although this research has mostly concentrated on frontal and temporal areas but not parietal parts of the brain. Based on our findings, we speculate that the right parietal areas implicated by our similarity analysis could be a left-hemisphere homologue area. Martin et al. (2022) hypothesize in their paper that the homologue areas might be “less specifically tuned for linguistic input” (page 366). If it is a general principle that homologue areas are less fine-tuned for detailed processing, this could explain why the right (but not the left) parietal areas show a high similarity in their oscillatory fingerprints between the language and memory task, potentially then reflecting concepts or representations, but not overly domain-specific content as one would find in pure language paradigms (e.g., Düzel et al., 2003; Spitzer et al., 2008).

### Between-task similarity dissociates between alpha and beta

Alpha and beta are often discussed together in the language (e.g., Cao et al., 2022; Piai et al., 2020) and memory literature (for review, see Fellner & Hanslmayr, 2017; Hanslmayr et al., 2016), alluding to the two rhythms substantiating the same function in both cases. However, in our analyses, alpha and beta behave divergently. While alpha shows a high similarity between tasks (but does not belong to a cluster), the beta range (19-26 Hz) belongs to a dissimilar cluster (Fig. 6B). This can partly be explained by motor beta band effects (discussed above), and potentially by “unspecific alpha” (discussed below). However, also functionally relevant areas such as temporal cortex (Fig. 6B) or, as we speculate, the homologue areas in the right parietal lobe (Fig. 6A) show distinct signatures between the tasks for alpha and beta, respectively. We thus wonder whether the treatment of the alpha/beta band as one phenomenon – as e.g., in the context of alpha/beta desynchronization effects – might need to be revisited. It is important to note again, that our results are, however, based on raw power and not power differences and that the present study is explorative in nature. Thus, future research could investigate this observed divergence between alpha and beta power in a hypothesis-driven manner and explore the link to the previously reported desynchronization effects. A replication of this finding would undoubtedly pose the question of functional importance of the two bands and whether either has a specific function in retrieval – a question we will explore for the alpha band in the following paragraph.

### Alpha similarity: specific to retrieval?

In the century since the first descriptions of the alpha rhythm (Adrian & Matthews, 1934; Berger, 1929), its functional relevance has been debated many times over (e.g., Fries, 2015; Hanslmayr et al., 2016; Jensen et al., 2002; Jensen & Mazaheri, 2010; Pfurtscheller et al., 1996; Toscani et al., 2010). Reasonably, we wonder about the origins of alpha and its similarity in our data: Can we assume a functional role when it comes to language and memory retrieval? In this context, it is important to note, again, that the alpha band itself is not part of the cluster substantiating that our impact measure is significantly different from zero (see Fig. 6A). However, in the similar regions (Fig. 6C), the difference in alpha (and beta) power between the tasks is small in some parts of the task windows (cf. for example right precuneus in Fig. 7A), pointing to a potential region-specific similarity in the alpha band that is not captured with our method. These findings make us wonder about a potential role of alpha when it comes to language and memory retrieval – or whether a less specific process might be underlying these results.

Above, we speculate that similarities between the tasks in the raw alpha power patterns could represent memory reinstatement or representations, based on previous studies, predominantly from the memory literature. Hanslmayr et al. (2016) theorize that a relative desynchronization in the alpha and beta band allows for the retrieval of memories through an “active engagement of cortical modules” (p. 19), which cannot be achieved with an overall high firing synchrony, potentially in an interplay with the hippocampus. An interpretational hurdle is created by the fact that most studies to date looked at alpha power decreases between conditions, while we are comparing raw alpha power between tasks. How both are related and whether they are indexing the same processes remains unknown; however, relative power decreases do not necessarily reflect an absence of this oscillation.

The alpha-beta decreases in the memory literature have recently been connected to eye movements (Popov & Staudigl, 2023), following a wider discussion of especially alpha in light of oculomotor activity (Liu et al., 2023; Pan et al., 2023; Popov et al., 2021). This challenges for example lateralized memory retrieval effects with visual hemifield stimulation during encoding (Waldhauser et al., 2016). Content-specific single trial results (Jafarpour et al., 2014; Michelmann et al., 2016; Staudigl et al., 2015) are harder to explain in this framework, although none of these studies have used eye tracking data for control analyses. Similarly, our study does not have eye tracking data either, an evident shortcoming, which prevents us from directly inspecting eye movements as a possible explanation for our results. We would like to note, however, that the spatial distribution of our similarity results does not fully fit the results of Popov and Staudigl (2023), which show a predominantly occipital distribution. Since the direct link between eye movements and alpha is not clear yet, it is hard to speculate what spatial profile we would expect across tasks. In a follow-up study that is currently underway, we added eye tracking to the protocol to investigate this potential explanation further.

Whether the alpha captured by our new analysis approach is a retrieval signature remains speculative and warrants further research. Overall, our language and memory retrieval alpha/beta signatures (cf. Fig. 4) are however congruent with earlier reported fingerprints for both tasks.

### Time is of the essence

Our analysis approach was inspired by an integrated approach of the time, frequency, and spatial features characterizing EEG data. We posed that shared functions can only be identified by their respective similarities across these dimensionalities. However, aligning the EEG signatures of two different tasks in time is no easy feat. We took guidance by within-task effects but must ultimately admit that we do not know how well this worked (cf. Fig. 6A). Crucially, we cannot deduct from diverging time courses that an underlying (sub-)process is different: that would assume that processes in different contexts always have the same time course. Given that cognitive processes show trial-to-trial (timing) variability in both behavior and associated neural activity (e.g., Chupina et al., 2025; Lundqvist et al., 2024) but also variability in different contexts (Carr et al., 2011; e.g., Ekman et al., 2017; Michelmann et al., 2018), this seems unlikely. A more fine-grained knowledge of serial and parallel organization of sub-processes (e.g., Dubarry et al., 2017; Marti & Dehaene, 2017) – and how they are reflected in the EEG signal – would not only afford for an easier comparison between tasks by making it possible to identify the neural signatures of to-be-compared sub-processes but also help in estimating trial-to-trial variabilities. To this end, the help of behavioural studies and targeted experimental manipulations is indispensable (Krakauer et al., 2017; Miller, 2010).

### Conclusions

In this paper, we present a new approach to studying similarities and dissimilarities between tasks using EEG data. We find that word and memory retrieval share oscillatory fingerprints in right parietal cortex but have dissimilar fingerprints in left temporal and frontal regions. We speculate that the right parietal areas could be left-hemisphere homologues, less tuned to specific memory or language functions yet involved in retrieval or representation of the relevant material.

## Acknowledgments

We thank Jan Mathijs Schoffelen for discussion of the data analysis approach. This study was funded by the Netherlands Organization for Scientific Research (Nederlandse Organisatie voor Wetenschappelijk Onderzoek [NWO], VI.Vidi.201.081 awarded to VP).

