## Supplementary material for "Shared and distinct oscillatory fingerprints underlying episodic memory and word retrieval"

#### Content

Table 1: Similar and dissimilar parcels 1
Figure 1: Source space results of impact analysis 3
Figure 2: Time-frequency task contrasts in regions of interest 4
Figure 3: Alpha power in task conditions 5

#### Supplementary Table 1

**Similar parcels, right hemisphere**

| | |
|---|---|
| *Parietal* | |
| Precuneus | Right Precuneus A31, area 31 (Lc1)<br>Right Precuneus A5m, medial area 5 (PEm) |
| IPL | Right Inferior Parietal Lobule A39c, caudal area 39 (PGp)<br>Right Inferior Parietal Lobule A39rd, rostrodorsal area 39 (Hip3) |
| *Frontal* | |
| Orbital G. | Right Orbital Gyrus A12/47l, lateral area 12/47 |

**Dissimilar parcels, left hemisphere**

| | |
|---|---|
| *Central* | |
| Precentral | Left Precentral Gyrus A4ul, area 4 (upper limb region)<br>Left Precentral Gyrus A6cdl, caudal dorsolateral area 6<br>Left Precentral Gyrus A4tl, area 4 (tongue and larynx region)<br>Left Precentral Gyrus A6cvl, caudal ventrolateral area 6 |
| Postcentral | Left Postcentral Gyrus A1/2/3tru, area1/2/3 (trunk region) |
| *Parietal* | |
| IPL | Left Inferior Parietal Lobule A40rv, rostroventral area 40 (PFop)<br>Left Inferior Parietal Lobule A40rd, rostrodorsal area 40 (PFt) |
| *Occipital* | |
| MVOC | Left Medioventral Occipital Cortex cCunG, caudal cuneus gyrus |
| | Left Medioventral Occipital Cortex rCunG, rostral cuneus gyrus |
| *Temporal* | |
| STG | Left Superior Temporal Gyrus TE1.0 and TE1.2 |

| | |
|---|---|
| | Left Superior Temporal Gyrus A22r, rostral area 22<br>Left Superior Temporal Gyrus A22c, caudal area 22<br>Left Superior Temporal Gyrus A41/42, area 41/42 |
| MTG | Left Middle Temporal Gyrus aSTS, anterior superior temporal sulcus<br>Left Middle Temporal Gyrus A21r, rostral area 21 |
| *Frontal* | |
| SFG | Left Superior Frontal Gyrus A6dl, dorsolateral area 6<br>Left Superior Frontal Gyrus A8dl, dorsolateral area 8 |
| MFG | Left Middle Frontal Gyrus A6vl, ventrolateral area 6<br>Left Middle Frontal Gyrus A8vl, ventrolateral area 8<br>Left Middle Frontal Gyrus IFJ, inferior frontal junction |
| IFG | Left Inferior Frontal Gyrus A44d, dorsal area 44<br>Left Inferior Frontal Gyrus A45c, caudal area 45 |

**Dissimilar parcels, right hemisphere**

| | |
|---|---|
| *Central* | |
| Precentral | Right Precentral Gyrus A4hf, area 4 (head and face region)<br>Right Precentral Gyrus A6cvl, caudal ventrolateral area 6<br>Right Precentral Gyrus A4tl, area 4 (tongue and larynx region) |
| Postcentral | Right Postcentral Gyrus A2, area 2<br>Right Postcentral Gyrus A1/2/3ulhf, area 1/2/3<br>Right Postcentral Gyrus A1/2/3tru, area1/2/3 |
| *Occipital* | |
| LOC | Right lateral Occipital Cortex msOccG, medial superior occipital gyrus |
| *Temporal* | |
| STG | Right Superior Temporal Gyrus A22c, caudal area 22<br>Right Superior Temporal Gyrus A38m, medial area 38 |
| MTG | Right Middle Temporal Gyrus A21r, rostral area 21 |
| *Frontal* | |
| SFG | Right Superior Frontal Gyrus A8dl, dorsolateral area 8 |
| MFG | Right Middle Frontal Gyrus A6vl, ventrolateral area 6<br>Right Middle Frontal Gyrus IFJ, inferior frontal junction |
| IFG | Right Inferior Frontal Gyrus A44op, opercular area 44 |

#### Supplementary Figure 1

### **Unmasked source space results of impact analysis**

Full results for the between-task impact analysis for brain regions, without statistical masking applied. Purple hues show dissimilar regions, brown hues show similar regions.

*Also see Figure 5E in the paper.*

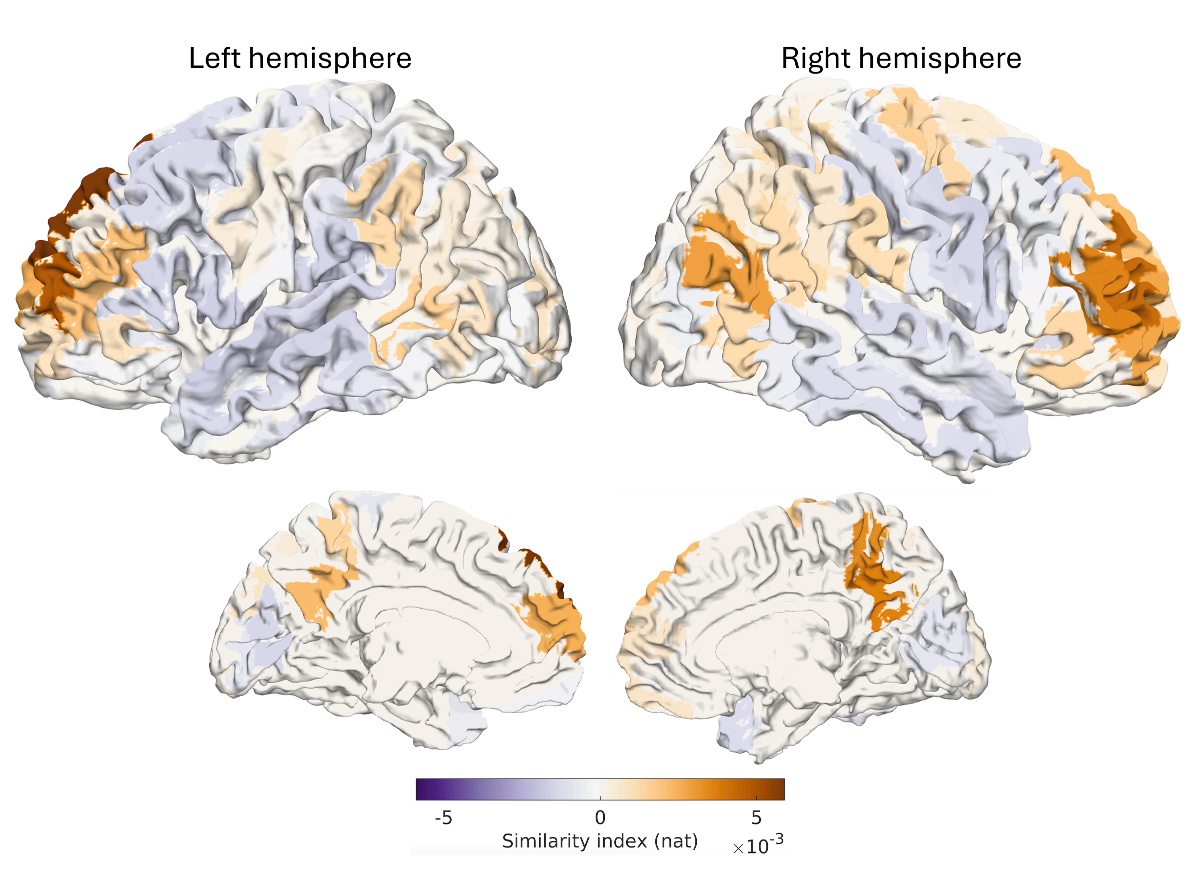

Supplementary Figure 2

#### Time-frequency representations in regions of interest, contrasting tasks

Contrasting time-resolved oscillatory power between the language and memory task (retrieval conditions only; *constrained* for the language task and *old* for the memory task). Red colors represent a higher absolute power in the language task, blue colors represent a higher absolute power in the memory task. Color limits are based on the maximum absolute power across both tasks in the given region of interest. Note the different time axes for the memory task (top) and the language task (bottom).
*Also see Figure 6 in the paper.*

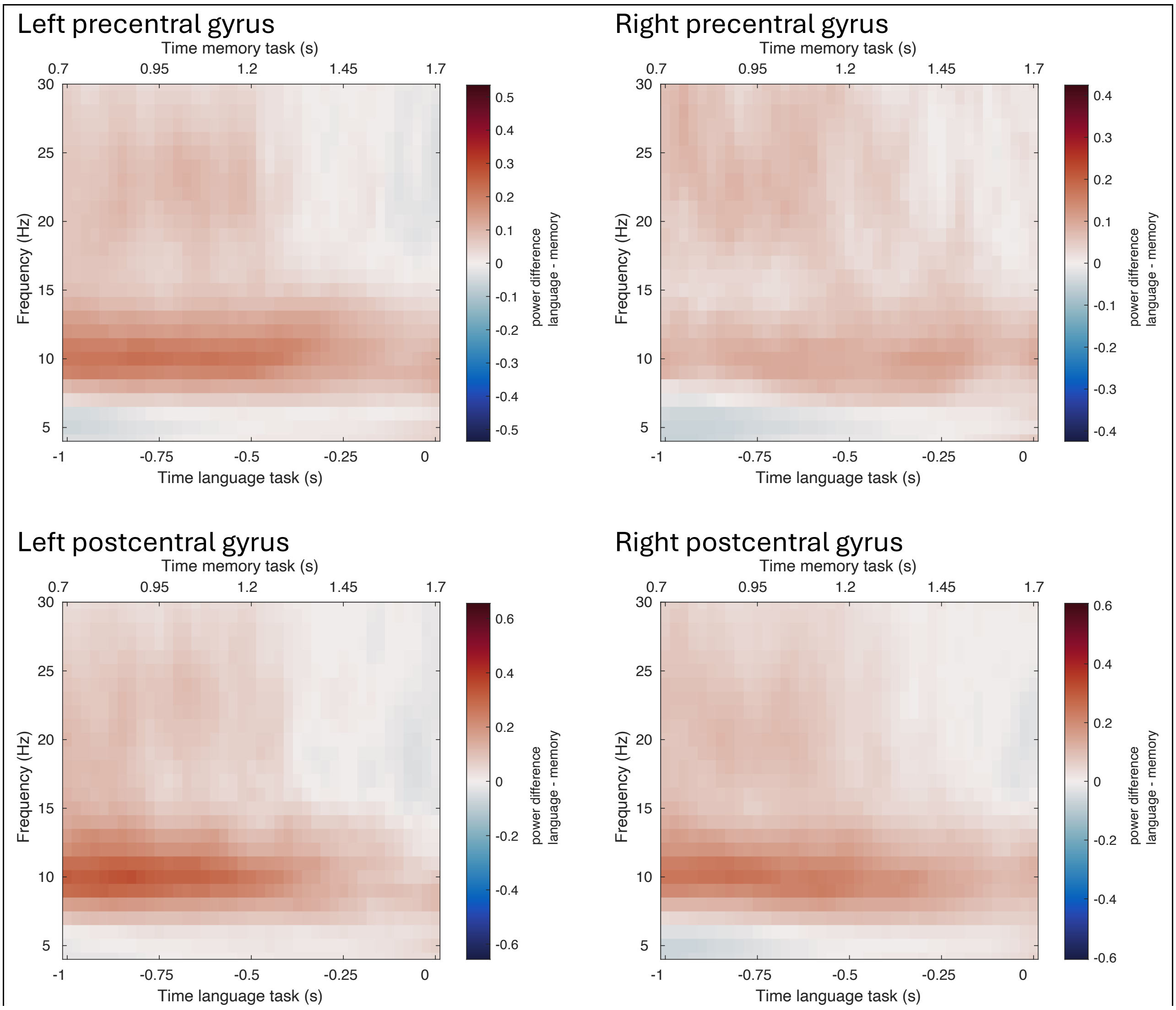

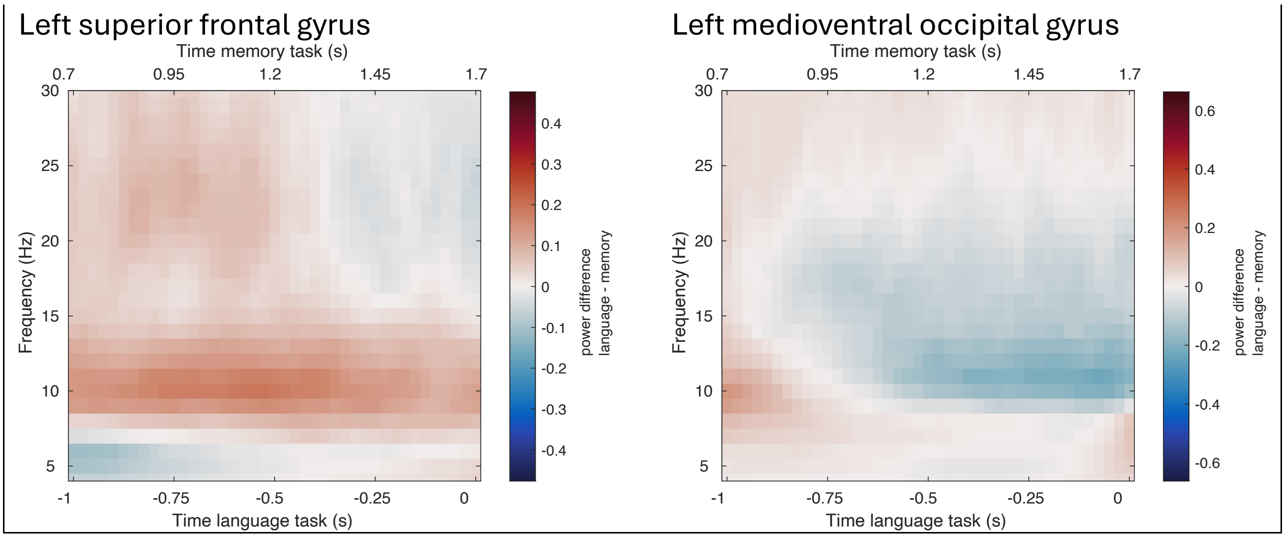


#### Supplementary Figure 3

#### **Alpha power in task conditions**

Alpha power across all electrodes per task and conditions, computed across the whole epoch (0.7 – 1.7 s after cue onset for the memory task; -1 – 0 s before picture onset for the language task).

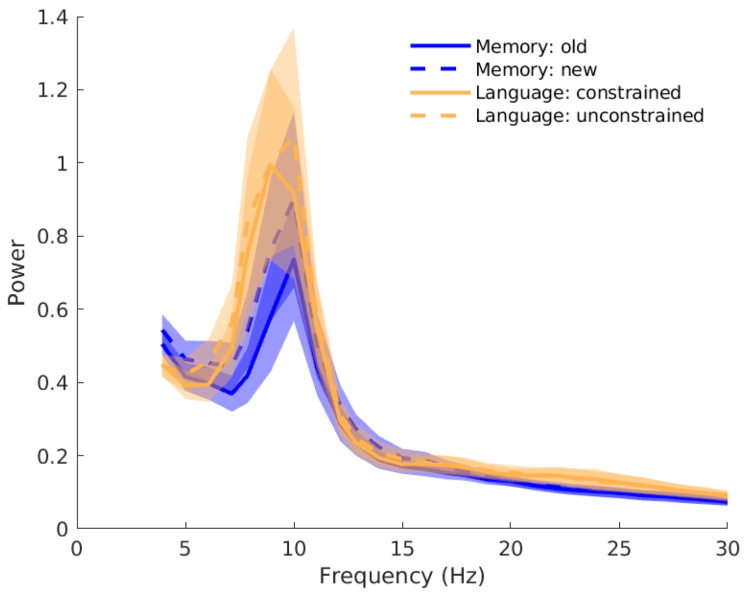
